# Why extinction estimates from extant phylogenies are so often zero

**DOI:** 10.1101/2021.01.04.425256

**Authors:** Stilianos Louca, Matthew W. Pennell

## Abstract

Time-calibrated phylogenies comprising only extant lineages are widely used to estimate historical speciation and extinction rates. Such extinction rate estimates have long been controversial as many phylogenetic studies report zero extinction in many taxa, a finding in conflict with the fossil record. To date, the causes of this widely observed discrepancy remain unresolved. Here we provide a novel and simple explanation for these “zero-inflated” extinction rate estimates, based on the recent discovery that there exist many alternative “congruent” diversification scenarios that cannot possibly be distinguished on the sole basis of extant timetrees. Consequently, estimation methods tend to converge to some scenario congruent to (i.e., statistically indistinguishable from) the true diversification scenario, but not necessarily to the true diversification scenario itself. This congruent scenario may in principle exhibit negative extinction rates, a biologically meaningless but mathematically feasible situation, in which case estimators will tend to hit and stick to the boundary estimate of zero extinction. To test this explanation, we estimated extinction rates using maximum likelihood for a set of simulated trees and for 121 empirical trees, while either allowing or preventing negative extinction rates. We find that the existence of congruence classes and imposed bounds on extinction rates can explain the zero-inflation of previous extinction rate estimates, even for large trees (1000 tips) and in the absence of any detectable model violations. Not only do our results likely resolve a long-standing mystery in phylogenetics, they demonstrate that model congruencies can have severe consequences in practice.

## Introduction

Understanding the historical dynamics of species formation and extinction is critical to explaining patterns of biodiversity (1, 2) and for quantifying the influence of environmental factors and human activity on the biosphere (3). The branching pattern of time-calibrated phylogenies of extant species (“extant timetrees”) is an important source of evidence, in addition to the fossil record, for reconstructing these dynamics; researchers thus often attempt to estimate speciation and extinction rates by fitting stochastic birth-death models to extant timetrees (4). Despite the widespread use of such phylogenetic approaches, there has been an enduring discrepancy that has never been satisfactorily explained: Analyses of extant timetrees using birth-death models often suggest that extinction rates were zero or negligible (5–11), contradicting the observation of high extinction rates in the fossil record and the fact that the majority of species ever to have existed are now extinct (2, 10). For example, Herrera (11) analyzed phylogenetic and fossil data of primates and observed that the phylogenetic data suggested negligible extinction rates while fossils showed that extinction rates were nearly equal to speciation rates.

Various explanations have been proposed for why extinction rates may be misestimated from phylogenies, all of which essentially invoke some type of model inadequacy (8, 12–15). However, the proposed mechanisms do not fully explain the observation that extinction rate estimates based on extant phylogenies are suspiciously often zero. Indeed, Rabosky (12) and Beaulieu *et al.* (14) showed that heterogeneity in rates across lineages (a violation of the assumption of homogeneity in birth-death models) often leads to an inflation — rather than deflation — of extinction rate estimates when these are obtained via birth-death model fitting (Fig. 3 therein). Similarly, Morlon *et al.* (13) proposed that failing to adequately account for variation in rates over time can lead to erroneous extinction rate estimates, although they never explained why this should specifically cause zero-inflated estimates. Purvis (8) suggested that biased species sampling or incompletely resolved phylogenies could lead to low extinction rate estimates. However, the mechanisms proposed by Purvis (8) are expected to alter extinction estimates by some factor, rather than completely erase any evidence for extinction. Importantly, it can be shown using simulations as well as empirical datasets (see below) that even when fitting birth-death models accounting for temporal variation, and even if trees are simulated under a birth-death model with homogenous rates across lineages and no sampling bias, extinction rate estimates are zero-inflated. A zero-inflation of extinction rate estimates even in the absence of any model violations can even be seen in the simulation results by Rabosky (12) and Beaulieu *et al.* (14), although these studies did not provide any explanation for that phenomenon. A zero-inflation of extinction rate estimates even without any model violations cannot possibly be explained by the above mechanisms. It is also notable that birth-death models are routinely fit to paleontological data without considering across-lineage heterogeneity and assuming a fairly simple model of time-variability (e.g., piecewise constant across different time-bins; 16) and yet typically estimate extinction rates to be similarly high as speciation rates (2).

Here we propose a novel and simple explanation for why extinction rate estimates from extant time-trees are often zero. Our explanation follows from our recent discovery that extant timetrees are generally insufficient for resolving historical diversification dynamics in the context of birth-death models (17). In contrast to other explanations, our reasoning holds true even in the absence of any model violations (i.e., when the data was indeed generated by a birth-death model and when the fitted model captures the variation in the data perfectly) and thus can explain zero-inflated extinction rate estimates in a much broader range of situations. Using simulations, we demonstrate that the common approach of selecting between alternative birth-death models using AIC does not resolve this issue; the best fit of a set of models frequently suggests zero extinction rate estimates even when in reality extinction was clearly non-zero. As we detail below, our explanation leads to multiple non-trivial predictions, which we fully confirm using simulated and empirical timetrees.

## Results and Discussion

### Model congruencies predicts zero-inflated extinction rate estimates

Our starting point is the general stochastic birth-death model with arbitrary time-dependent speciation rate *λ*, time-dependent extinction rate *μ* and random lineage sampling at present day with sampling fraction *ρ* (13, 17). Note that here we use “diversification scenario” to refer to a single specific choice of *λ* and *μ* over time, as well as a specific value of *ρ*. We recently showed that historical diversification dynamics cannot be unambiguously reconstructed from extant timetrees alone (17): for any given diversification scenario there exist many alternative (“congruent”) scenarios that would generate extant timetrees with the same probability distribution as the proposed scenario. Such congruent scenarios exhibit identical likelihood functions and cannot possibly be distinguished using extant timetrees alone, no matter how large and complete. The set of all congruent scenarios, called a “congruence class”, is infinitely large, infinite dimensional, and contains a myriad of similarly plausible and yet markedly different scenarios. Crucially, when fitting birth-death models to an extant timetree, either via maximum-likelihood or using Bayesian inference, estimators will generally converge towards the congruence class of the true diversification scenario but not necessarily to the true diversification scenario itself.

The above realization suggests that in practice extinction rate estimates obtained entirely from extant timetrees (no matter how large) will often be very wrong (17, 18). The question is: why are these (wrong) extinction rate estimates often exactly zero, instead of simply “random” positive numbers? In other words, why is the distribution of extinction rate estimates seen in the literature zero-inflated? We propose that the answer to this mystery stems from the fact that the likelihood function of a birth-death model is mathematically well-defined even for extinction rates that are negative during some or all time points (19). If one were to consider models with partly or fully negative extinction rates, then the corresponding congruence class of the true historical diversification history, denoted ℍ, would include many additional scenarios with partly or fully negative *μ* and yet positive *λ* (see for example Supplement S1.4 in (17) for constructing such congruent scenarios). Of course, a scenario with *μ* < 0 is biologically meaningless, but if one were to permit such scenarios, it could often be the case that such a scenario is “closest” to ℍ (i.e., has the highest likelihood) among the set of considered scenarios, even compared to those scenarios with positive *μ*. This is not at all paradoxical once one recognizes that what one is really estimating is the congruence class of the true diversification history and not the true diversification history itself and that any scenario with negative *μ* is congruent to a myriad of scenarios with positive biologically plausible *μ*. However, imposing the biologically motivated constraint that *μ* ≥ 0 places a boundary in parameter space that likelihood-optimizers and Bayesian samplers will tend to run up against, thus yielding estimates for *μ* that are partly or entirely zero (illustration in Fig. 1A). Such situations can in principle occur for birth-death models of arbitrary complexity, for example with time-dependent *λ* and *μ* (as we demonstrate below). Note that whether the fitted model exhibits negative extinction rates (if allowed) ultimately depends on the full set of scenarios considered (e.g., the specific functional forms fitted).

**Figure 1:**
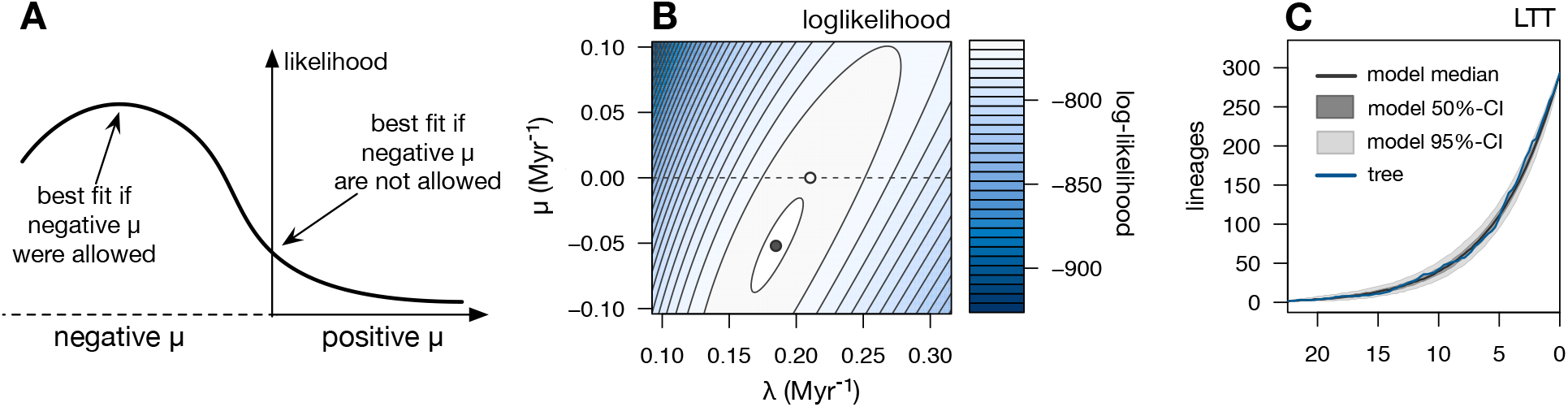
Conceptual illustration. (A) For any given extant timetree, the likelihood function will generally be maximized in a region of parameter space close to the congruence class of the true diversification history (including scenarios with partly or fully negative *μ*), but not necessarily close to the true diversification history itself. This maximum may even be located in a region where *μ*; < 0. Constraining *μ* to be positive may merely result in a “compromised” fit where *μ* = 0. In the above illustration we focus on *μ* as a single fitted parameter for simplicity, however in practice fitted models may depend on multiple parameters, for example if *μ* and *λ* have complex functional forms. (B) Contour-plot of the log-likelihood of an empirical timetree of 293 Trochilidae species (20), under constant-rate birth-death models with various speciation rates (horizontal axis) and extinction rates (vertical axis), and with fixed known sampling proportion. When extinction rates are allowed to be negative, the maximum-likelihood scenario exhibits a negative extinction rate (black dot). Constraining *μ* to non-negative values will yield a maximum-likelihood estimate that is zero (white dot). The dashed horizontal line at the *μ* = 0 boundary is shown for reference. (C) Lineages-through-time (LTT) curve of the Trochilidae tree (blue curve), compared to LTTs generated by the fitted constant-rates birth-death model (black curve shows median, shades show 50% and 95% confidence intervals. Observe the good agreement between the model’s and the tree’s LTTs. The fitted model adequately describes the tree; it cannot be rejected using parameteric bootstrapping based on the Sackin (21) and Colless (22) tree imbalance statistics, the distribution of node ages and the distribution of branch lengths.

Figure 1B–C illustrates the above reasoning using an empirical timetree of 293 Trochilidae (hummingbird) species (20). If we only consider scenarios where speciation and extinction rates are constant through time, the likelihood has its global maximum at a scenario with a negative extinction rate (Fig. 1B). As expected, fitting constant-rates birth-death models to the Trochilidae tree yields a negative extinction rate when *μ* is allowed to be negative and a zero extinction rate when *μ* is constrained to non-negative values (Fig. 1B). Importantly, in this case the fitted model with zero extinction rate actually explains the data very well (Fig. 1C), and could not be rejected based on any model adequacy test considered (using parametric bootstrapping and considering the Sackin (21) and Colless (22) tree imbalance statistics, the distribution of edge lengths based on a Kolmogorov-Smirnov test, and the distribution of node ages based on a Kolmogorov-Smirnov test; P>0.05 in all cases). Hence, the fact that *μ* is estimated to be zero is unlikely due to an inadequacy of the model to explain the data at hand; rather, the maximum-likelihood scenario with negative *μ* is probably congruent to another (biologically plausible) diversification scenario close to the true (but unknown) hummingbird diversification history.

Based on the above reasoning, we can make the following testable predictions for extinction rate estimates obtained via maximum likelihood estimation (we do not consider Bayesian estimation in this paper, although similar arguments would apply). First, erroneously obtaining zero extinction rate estimates should be common even in situations without any detectable model violations, i.e., even where the fitted models explain the data well. Second, in almost all cases where the extinction rate is erroneously estimated to be zero at one or more time points, one should obtain a negative extinction rate estimate at one or more time points if one were to allow for such estimates. In particular, when allowing negative extinction rates, the distribution of estimated extinction rates should no longer be zero-inflated. Third, estimating a negative extinction rate (if allowed) should increase the chances of obtaining a zero extinction rate estimate when constrained to non-negative values (e.g., due to the optimization routine or sampler getting “trapped” on the *μ* = 0 boundary). Note, however, that this is not a strict requirement, i.e., in some cases the estimated *μ* might be negative if allowed and yet strictly positive if constrained, for example if the likelihood function has multiple local maxima. Fourth, in cases where the estimated extinction rate is positive even if allowed to be negative, this estimate should typically be similar to the estimate obtained when constraining *μ* to be non-negative; this is again not a strict requirement due to the possibility of multiple local maxima and the fact that the optimal value for one parameter generally depends on the values of the other parameters. Fifth, fixing the speciation rate to its true value during fitting (thus, “collapsing” the congruence class to a single scenario) should yield much more accurate extinction rate estimates and should eliminate their zero-inflation. Sixth, even for fitted models with negative extinction rates, these models should be close to the true diversification history’s congruence class (provided, of course, sufficient data for fitting and a sufficiently flexible model). As we describe below, we confirmed all six predictions using numerical simulations, and also confirmed predictions 2–4 using 121 previously published timetrees from across the Eukaryotic Tree of Life.

### Analysis of simulated data

To test our predictions, we simulated 500 extant timetrees using birth-death scenarios with known *λ* and *μ*, and then fitted models while either constraining *μ* to be non-negative or allowing *μ* to also attain negative values (*λ* was always constrained to be non-negative, and *ρ* was fixed to its true value). To underscore the fact that the underestimation of *μ* is not simply due to stochasticity stemming from small data sets, we considered large timetrees comprising 1000 tips. For simplicity, in all cases we focused on the extinction rate at present-day (henceforth denoted *μ_o_*), although all conclusions would apply similarly to any other time point. The *λ* and *μ* used in every simulation were time-dependent and their profiles defined according to simple stochastic processes (of Ornstein-Uhlenbeck type; 23). This approach was chosen in order to obtain a large variety of realistically complex diversification dynamics, as typically observed in the fossil record (24). The present-day *μ_o_* was chosen randomly and uniformly between 0.1 · *λ_o_* and *λ_o_*, and at all times *μ* was constrained to between 0.1 · *λ_o_* and *λ_o_*; hence, the extinction rate was clearly non-zero. We then used maximum-likelihood to fit various functional forms for *λ* and *μ* commonly used in the literature (25): *λ* was either i) assumed to be constant over time, ii) assumed to vary exponentially (*λ*(*t*) = *αe^βt^*), iii) assumed to vary exponentially plus a constant (*λ* = *αe^βt^* + *γ*), or iv) assumed to vary linearly (*λ*(*t*) = *λ_o_* + *αt*). Similar profiles were considered for *μ*, thus yielding 4×4 alternative combinations. We refer to these 16 models as “ELC” models, for Exponential/Linear/Constant; depending on whether we constrain *μ* to be non-negative or allow *μ* to be negative, we call a fitted model “constrained” or “unconstrained”, respectively. For every tree we fit all 16 ELC models and then selected the best fitted model using AIC, following common practice (13). It is worth noting that real diversification dynamics emerge from a myriad of complex biological/environmental processes that will probably never be exactly captured by any human-made mathematical model; instead, the typical objective is to find an approximate model that adequately describes the data at hand. To reduce errors in the estimated extinction rates stemming from poor model fits (26, 27) we only considered simulations where the best fitted model adequately described the data, as evaluated using parametric bootstrapping with a Kolmogorov-Smirnov test comparing the distribution of node ages in the tree versus the fitted model (380 out of 500 simulations were kept). For every such simulation, we examined whether the estimated present-day extinction rate in the best constrained fitted model (denoted 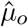) was zero. To each tree, we also fit an unconstrained version of the best model — i.e., using the same functional forms for *λ* and *μ* but allowing *μ* to be negative, thus obtaining a potentially new estimate for the present-day extinction rate (denoted 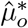).

For all considered simulated trees the best fitted constrained as well as unconstrained model closely matched the deterministic LTT of the true diversification scenario used in the simulation as well as the LTT of the simulated tree itself (fraction of explained variance *R*^2^ ≥ 0.99 in all cases, Supplemental Fig. S1B), confirming that the fits were successful (examples in Supplemental Fig. S2). Despite the good fits and despite the large size of the input trees, present-day extinction rate estimates 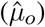 poorly reflected the true extinction rates used in the simulations (Fig. 2A). This is consistent with our general argument that extinction rates cannot be reliably estimated from extant timetrees in the absence of further information, due to fundamental identifiability limits (17). Importantly, for 93 out of 380 simulated trees with sufficiently good fit (24.5%), the best supported constrained model yielded a present-day extinction rate estimate of exactly zero (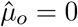, Fig. 2A and Fig. 3A). A zero 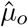 is in conflict with the fact that the true extinction rate was at all times clearly non-zero (*μ* ∈ [0.1 · *λ_o_, λ_o_*]). This fraction of zero extinction rate estimates is also much larger than what would be expected if extinction rate estimates were randomly and continuously distributed; in other words, the distribution of extinction rate estimates is zero-inflated, consistent with observations in the literature and with our predictions. Moreover, we did not observe any strong nor significant positive correlation between the goodness-of-fit (in terms of the deterministic LTT’s *R*^2^, or in terms of the P-value of the KolmogorovSmirnov test for node ages) on the one hand and the relative estimation error 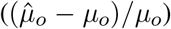 on the other hand (*P* > 0.05 and Spearman’s *ρ* < 0.1, Supplemental Fig. S1), suggesting that zero-inflated extinction estimates are unlikely the result of poor model fits. In all cases where 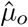 was zero, the unconstrained variant yielded a negative 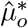, again consistent with our predictions (Fig. 2C). Reciprocally, for nearly all trees with negative 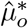 we obtained a zero 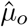, again consistent with our predictions (Fig. 2C). In particular, and in contrast to 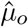, the distribution of 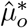 was not zero-inflated (Fig. 3B). For trees where the fitted 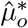 was positive, the fitted 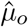 was almost always identical to 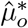 (up to numerical accuracy), again confirming our predictions (Fig. 2C). These results confirm our interpretation that the zero-inflation of extinction rate estimates stems from “cutting off” extinction rate estimates that would otherwise be negative (Fig. 2B, example in Fig. 4).

**Figure 2:**
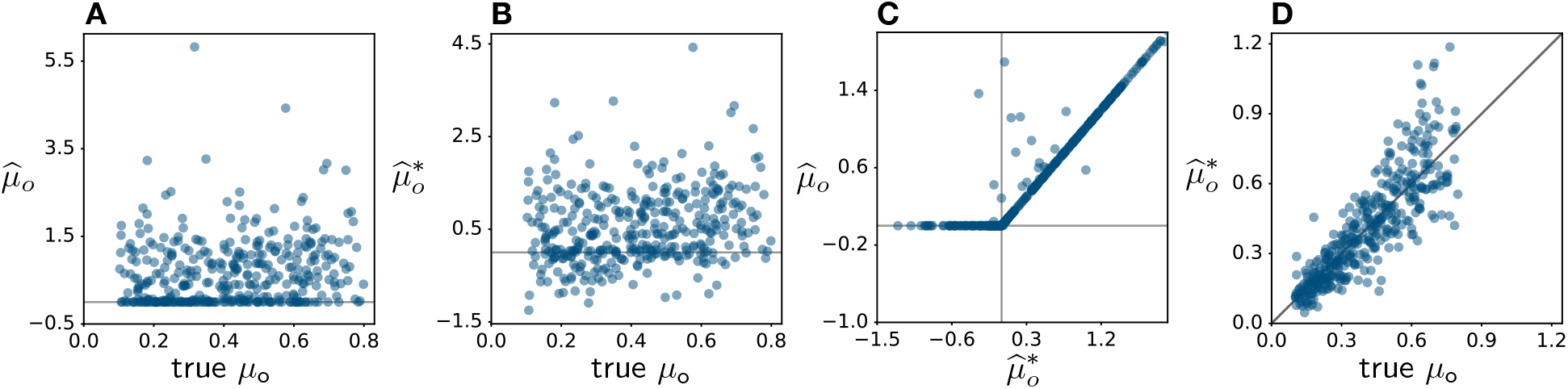
Testing predictions using simulations. (A) Present-day extinction rates fitted to 500 extant timetrees simulated under birth-death models with time-dependent *λ* and *μ*, while requiring the fitted extinction rate to be non-negative (one point per tree). Various commonly used models (“ELC” models, described in the main text) were fitted, and the best fitted model was selected based on AIC. Vertical axis: fitted present-day extinction rate 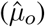. Horizontal axis: true present-day extinction rates (*μ_o_*). (B) Present-day extinction rates fitted to the same trees and with the same best models as in A, but allowing for negative extinction rates. (C) Present-day extinction rates fitted while allowing negative extinction rates 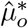, horizontal axis) compared to the case where negative rates are not permitted 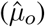, for the same trees as in A (one point per tree). (D) Present-day extinction rates 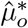 (vertical axis) fitted while fixing the speciation rate *λ* to its true profile, compared to the true present-day extinction rate (horizontal axis). The diagonal is shown for reference. All simulated trees comprised a large number of tips (1000), to avoid estimation error due to small data sizes. All rates are expressed in Myr^−1^. For histograms of the rate estimates see Fig. 3.

**Figure 3:**
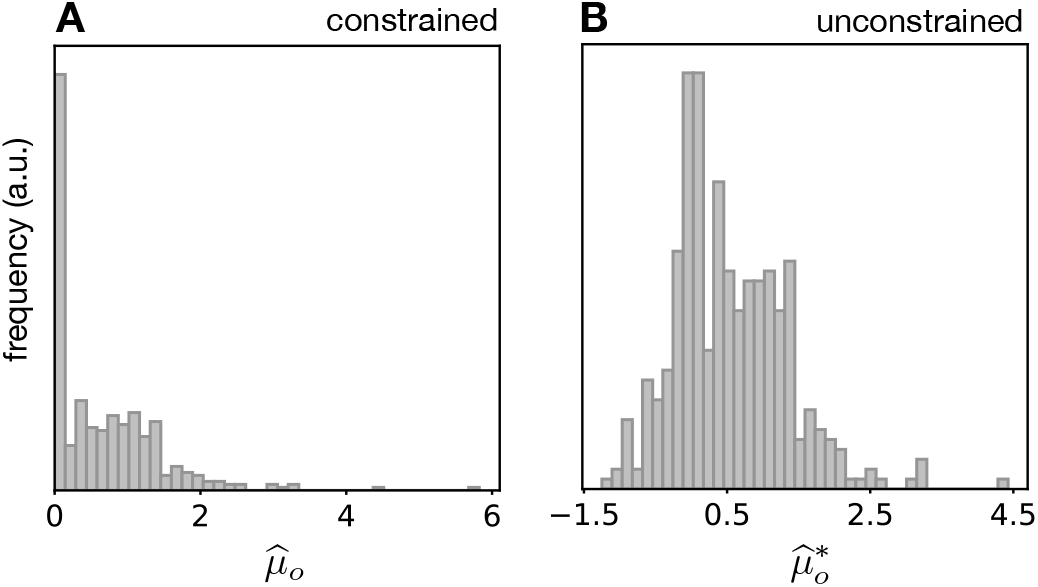
Distribution of estimated 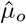 and 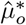. (A) Histogram of present-day extinction rates 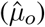 estimated for 380 simulated timetrees (as in Fig. 2) based on the best fitted ELC model, while constraining extinction rates to be non-negative. Observe the clear zero-inflation of present-day extinction rate estimates. (B) Histogram of estimated present-day extinction rate 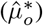 across the same timetrees as in (A), based on the same fitted ELC models as in (A) but allowing for negative extinction rates.

**Figure 4:**
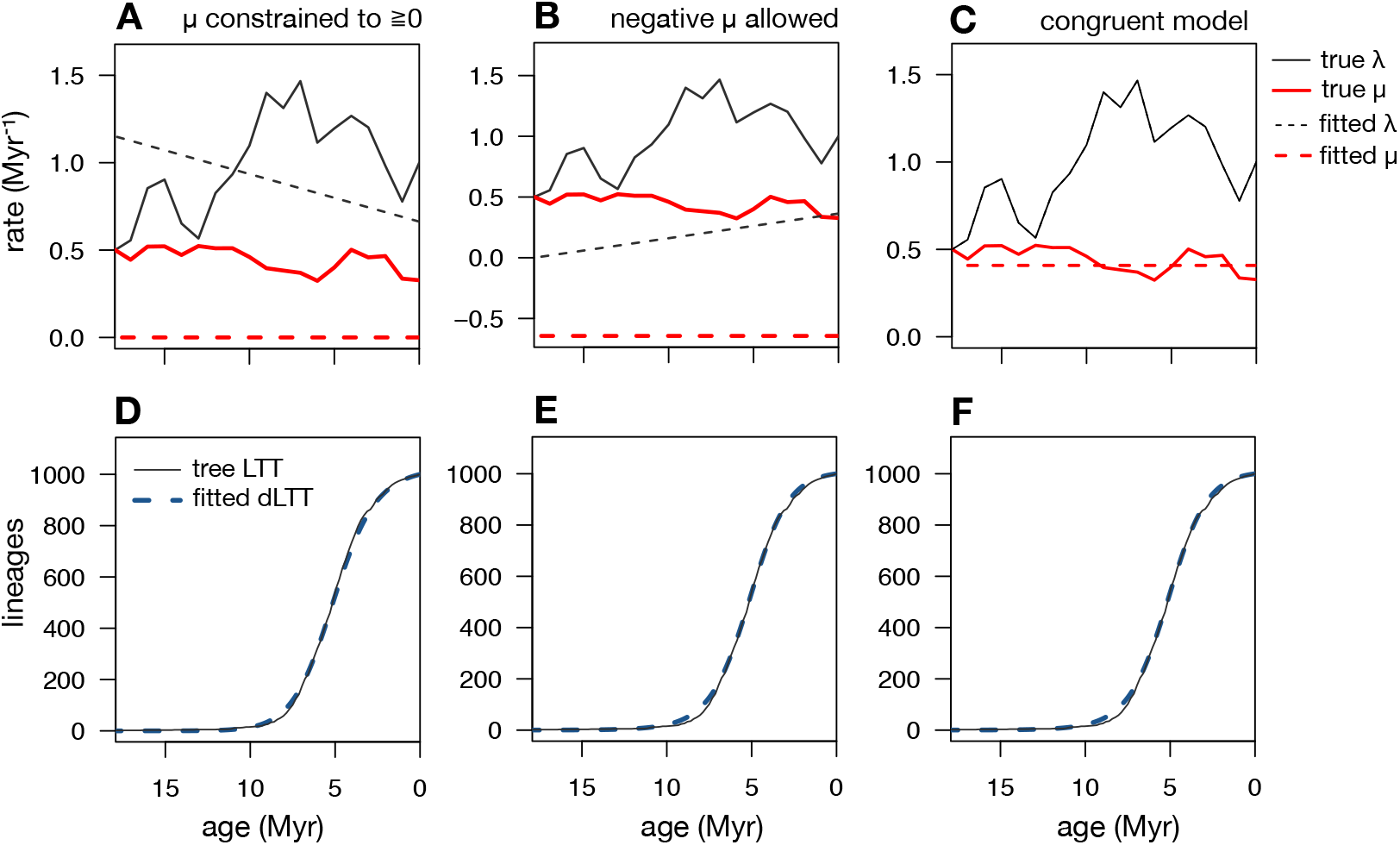
Fitted models are approximately congruent to the true diversification scenarios. Example ELC model fit (and selected according to AIC) to a simulated timetree with time-variable speciation rates (*λ*) and extinction rates (*μ*). (A) True *λ* and *μ* (continuous curves) compared to fitted *λ* and *μ* (dashed curves), when *μ* is constrained to be non-negative. (B) True *λ* and *μ* (continuous curves) for the same tree as in A, compared to *λ* and *μ* (dashed curves) fitted when *μ* is allowed to be negative. The negative fitted 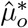 suggests that in A the zero 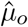 was simply “cut-off” at that boundary. (C) Diversification scenario congruent to the model fitted in B, but exhibiting the correct extinction rate; observe that this scenario is close to the true scenario, which means that the congruence class of the model fitted in B is close to the congruence class of the true scenario. (D–F) Lineages-through-time (LTT) curve of the simulated tree (continuous curves) compared to the deterministic LTTs of the fitted models shown in A–C. The good match between the tree’s LTT and the fitted models’ deterministic LTTs is consistent with the conclusion that the fitted models are close to the congruence class of the true scenario. For additional examples see Supplemental Fig. S3.

The above analyses reveal that the zero extinction rate estimates in our simulations resulted from a restriction to non-negative rates in cases where the maximum-likelihood fit would actually be negative (if allowed). Note that the frequent occurrence of likelihood peaks in biologically implausible regions of model space (e.g., Fig. 1B), despite the large tree sizes, makes complete sense in light of model congruencies: When fitting birth-death models one is at most estimating the congruence class of the true diversification history, rather than the true diversification history itself, and this congruence class typically includes a myriad of scenarios with negative extinction rates. For example, in the common case where specific parameterized functional forms are fitted for *λ* and *μ* (as is done here), one will obtain a combination of parameters whose corresponding *λ* and *μ* curves come closest to some member of the true diversification scenario’s congruence class, even if the true diversification scenario was very different. This interpretation is supported by the observation that for all considered simulations, the fitted model indeed closely matched the true scenario’s deterministic LTT (dLTT, which fully determines the congruence class), even in cases where 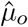 was zero (Supplemental Fig. S2). To further test this interpretation, for each best fitted unconstrained model with negative 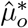, we determined the unique congruent diversification scenario with the correct *μ*, that is, the member of the fitted model’s congruence class that happens to have the correct *μ*. As expected, this congruent scenario typically had a speciation rate profile close to the truth, confirming our interpretation that the fitted models, even those with negative 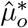, are actually close to the true diversification scenario’s congruence class (examples in Fig. 4C and Supplemental Fig. S3). Similarly, when we re-fitted ELC models to the trees while fixing *λ* to its true profile, the extinction rate estimates were much more accurate and no longer zero-inflated (in fact, none of them were zero, Fig. 2D), again consistent with our predictions.

The above results cannot be attributed to model-inadequacy, i.e., to the possibility that ELC models are not flexible enough to adequately approximate the true profiles of *λ* and *μ* over time used in the simulations, for three reasons. First, there is no reason to expect that this type of model inadequacy should lead to zero-inflated extinction rates, instead of simply a mix of over- and under-estimation errors. Second, and more importantly, in all considered simulations the best fitted constrained model matched the data well, based on a Kolmogorov-Smirnov test on the distribution of node ages and based on the fact that each tree’s LTT (which contains all information in the context of birth-death models; (17, 28, 29)) was completely or nearly completely contained within the 95% confidence interval of LTTs generated by the best fitted model. Similarly, the deterministic LTTs (dLTTs) of the best fitted models closely resembled the dLTTs of the true scenarios used in the simulations (*R*^2^ > 0.99 in all cases, examples in Supplemental Fig. S2). Since the dLTT of a birth-death model fully determines its congruence class (17) and the probability distribution of generated trees (when *μ* is non-negative), this means that the best fitted constrained and unconstrained models indeed converged towards the true congruence class and that the best fitted constrained models would generate trees similar to those of the true diversification scenario; this pattern is fully consistent with our theory and rules out the possibility of serious model inadequacies. Third, as mentioned earlier, there was no significant positive correlation between the considered goodness of fit measures and the relative estimation error 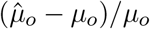, Supplemental Figs. S1A,B), contrary to what would be expected if model inadequacy caused a deflation of extinction rate estimates. Similarly, we did not observe a negative correlation between the considered goodness of fit measures and the absolute relative estimation error 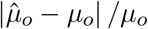, Supplemental Figs. S1C,D), contrary to what would be expected if model inadequacy was the cause of erroneous (in any direction) extinction rate estimates.

### Analysis of empirical timetrees

Next, to test the relevance of our interpretation to real datasets, we examined 121 previously published time-trees, spanning the eukaryotic Tree of Life. The size of the trees ranged from 13 to 11,638 species, and their root age ranged from 6.6 to 431 Myr. To facilitate comparison with our simulation results, we fit the same ELC models as described before and chose the best model based on AIC. The sampling fraction *ρ* was estimated separately based on the literature and fixed during model fitting (references and details in Supplemental Data File 1). In each case, we either constrained *μ* to be non-negative or allowed *μ* to be negative, and compared the obtained present-day extinction rate estimates with or without the constraint 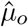 and 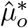, respectively). To further avoid biases due to bad model fits, we omitted 8 trees for which the best fitted model was rejected based on the distribution of node ages and using a Kolmogorov-Smirnov test (P<0.05).

Similar to our simulations, for all 113 considered empirical trees the fitted models explained the data well, with each tree’s LTT falling entirely or almost entirely within the 95% confidence interval of LTTs generated by the best fitted constrained model (Supplemental Fig. S4), and with the model’s dLTT capturing nearly all of the variance of the tree’s LTT (*R*^2^ > 0.99, Supplemental Fig. S5E). For 37 out of 113 trees the fitted model exhibited a present-day extinction rate of exactly zero 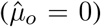; this high fraction of zero extinction rate estimates is consistent with the zero-inflated estimates commonly seen in the literature and in our simulations. In all cases where 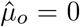, the fitted 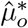 was negative, whereas a positive 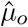 nearly always coincided with a positive and identical 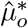 (Fig. 5A), consistent with our predictions and our simulation results. We mention that, as in our simulations, it is easy to construct models congruent to the fitted models (even those with negative 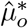) with plausible non-negative *λ* and *μ*; some of these congruent models are likely close to the taxon’s true diversification history (17), although in contrast to our simulations it is impossible to know which ones those are.

**Figure 5:**
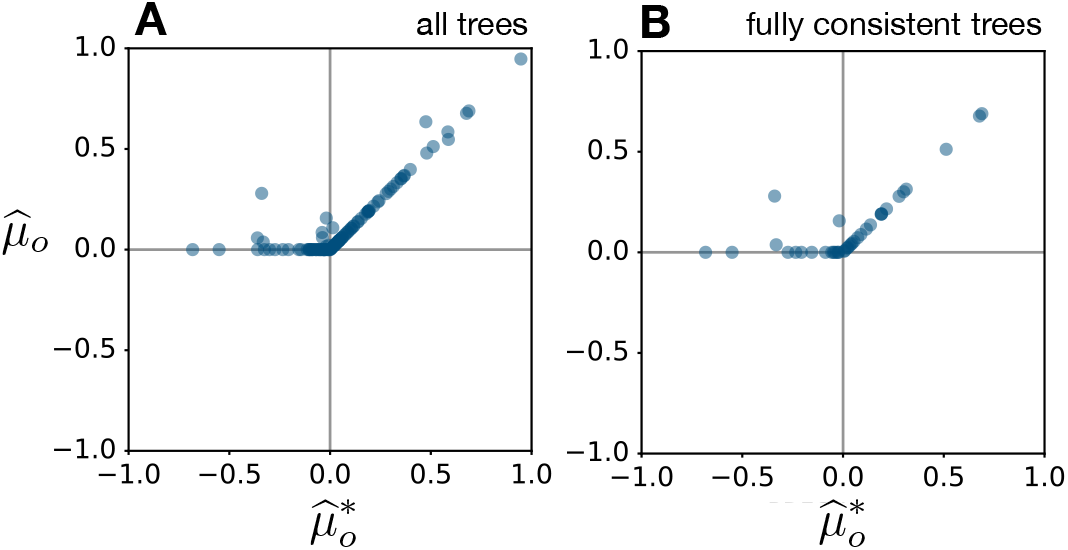
Testing predictions with real data. Present-day extinction rates estimated from empirical extant timetrees of various taxa (one point per tree), by fitting 9 commonly used models (“EC” models, explained in the main text) and choosing the best model via AIC. Horizontal axes show the fitted extinction rate while allowing for negative extinction rates 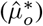, vertical axes show the fitted extinction rate when negative rates are not allowed 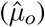. All units are in Myr^−1^. (A) Showing all trees where the best fitted model adequately explains the distribution of node ages (using a Kolmogorov-Smirnov test). (B) Showing only trees where the best fitted model also adequately explains the distribution of edge lengths (using a Kolmogorov-Smirnov test), the tree’s Sacking and Colless imbalance statistics. Only trees with a fitted 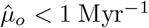 are shown in order to focus on near-zero and negative rate estimates.

As we discussed above, erroneous extinction estimates have previously been attributed to unmodeled variation in diversification rates *between* different lineages (12, 15) existing at any given time point. While previous research has not identified any mechanism by which such model inadequacy should generally lead to zero-inflated rate estimates, we nevertheless consider this possibility here. Specifically, for every tree we checked to what extent the best fitted model adequately explained not only variation through time (captured by the LTT) but also variation across lineages using parametric bootstrapping and four test statistics: the Sackin and Colless (22) statistics, the distribution of edge lengths (using a Kolmogorov-Smirnov test), and the distribution of node ages (using a Kolmogorov-Smirnov test). Perhaps not surprisingly, a large fraction of models were rejected based on at least one of these tests, with only 40 out of 113 trees passing all tests — that is to say, many of the empirical datasets were not fully consistent with the fitted homogeneous birth-death models. Indeed, the majority of rejected models were rejected based on the basis of the Sackin and/or Colless test statistics, which are specifically designed to detect tree imbalances, for example caused by rate heterogeneities across lineages. For 13 of the 40 remaining trees, the best fitted models exhibited a zero extinction rate, which corresponds to a similar fraction as that observed in the larger tree set. Moreover, we did not observe any significant positive correlation between the statistical significance (P-value) of any of the deployed adequacy tests and 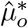, which one would expect if model inadequacies were the main cause of deflated extinction rate estimates (Spearman’s rank correlations were either negative, or non-significant, see Supplemental Fig. S5). Hence, processes leading to deviations from the assumptions of birth-death models do not seem to provide a plausible alternative explanation for the high frequency of zero (or negative, if allowed) extinction rate estimates in these empirical trees.

## Conclusions

The existence of vast stretches of congruent (i.e., statistically indistinguishable) and yet markedly different diversification scenarios severely compromises the reliability of extinction rate estimates based entirely on extant timetrees (17). However, even after our previous discovery (17), it was unclear whether and how model congruencies could explain the zero-inflated extinction rate estimates from extant timetrees widely observed in the literature. Here we have shown that these zero extinction estimates are probably due, at least in part, to the fact that numerical estimators will converge to the closest models in the congruence class of the true but unknown diversification history and not necessarily to the true diversification history itself — even if the closest models are biologically meaningless. Importantly, this can occur even if the data are well-described by the model (or more precisely, by its congruence class), that is, even in the absence of detectable model violations. It is of course possible that previously proposed mechanisms due to model inadequacy or misspecification (8, 12, 15) also lead to erroneous extinction rate estimates in real datasets. However, none of these mechanisms has been shown to explain the zero-inflation of extinction estimates. In fact, our observation that the proportion of zero extinction estimates was similar for those empirical trees that passed all four of our model adequacy tests versus those that didn’t, suggests that model inadequacy is not an important additional cause of zero-inflated extinction estimates in real datasets. Regardless of whether our proposed mechanism is the sole or even main cause of zero-inflated extinction estimates, our results clearly show that highly erroneous zero-inflated extinction estimates occur even in situations without any detectable model inadequacy and even for large trees with thousands of tips. Our results lead us to concur with other researchers that many previous estimates of extinction rates from extant timetrees alone are not reliable (12), and that most zero extinction rate estimates from such studies are almost certainly wrong (2). More generally, our findings provide a clear case in point that birth-death model congruencies have likely been seriously confounding macroevolutionary studies for decades.

## Materials and Methods

All simulations and maximum-likelihood fitting of birth-death models were performed using the R package castor v1.6.5 (30). All simulated trees had 1000 tips. All times are measured in Myr, and rates are measured in Myr^−1^. To simulate a variety of diversification scenarios we generated random profiles for *λ* and *μ* according to independent Ornstein-Uhlenbeck (OU) stochastic processes (23). This approach was chosen in order to cover a wide range of diversification scenarios with realistic temporal complexity. For *λ*, the present-day value *λ_o_* (serving as initial condition for the OU process) was set to 1 Myr^−1^, the stationary expectation was set to *λ_o_*, the stationary standard deviation was set to 0.5 · *λ_o_* and the relaxation rate was set to 0.1 Myr^−1^. For *μ*, the present-day value *μ_o_* was chosen randomly between 0.1 · *λ_o_* and 0.8 · *λ_o_*, the stationary expectation was set to *μ_o_*, the stationary standard deviation was set to 0.5 · *μ_o_* and the relaxation rate was set to 0.1 Myr^−1^. Both *λ* and *μ* were specified on a discrete time grid spanning 100 Myr and having a time-step of 1 Myr, according to the exact distribution of OU paths, with the exception that values below 0.1 Myr^−1^ were replaced with 0.1 Myr^−1^ to ensure that speciation and extinction rates were always above a detectable threshold. The sampling fraction *ρ* was chosen randomly and uniformly on a logarithmic scale from 0.01 to 1. For every given profile for *λ* and *μ* and chosen *ρ*, we simulated an extant timetree using the castor function `generate_tree_hbd_reverse` (31). We then fitted ELC models to the tree using the castor function fit_hbd_model_parametric with appropriately defined functional forms for each ELC model (see main text for functional forms) and with options `“condition=’auto’, relative_dt=1e-3, max_start_attempts=100, fit_control=list(eval.max=10000, iter.max=1000, rel.tol=1e-12, step.min=0.0001)`”. Fitting was agnostic of the simulation parameters, i.e., as if no further information was available apart from the tree itself and the sampling fraction *ρ* (which was fixed to its true value). As a first guess for the parameters (option `“param_guess”`) we set the exponent (in the case of exponentially varying rates) to zero, and the present-day rates to values obtained by first fitting a constant-rates BD model with known *ρ*. The bounds or the fitted parameters were chosen such that *μ* was necessarily non-negative. After fitting all ELC models to a tree, we chose the ELC model that had the smallest AIC, following common practice. We then re-fitted that model with parameter bounds adjusted to allow for negative *μ*, and compared the estimated present-day *μ_o_* obtained with and without constraining *μ* to non-negative values (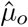 and 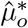, respectively). To also examine how the extinction rate estimates change if *λ* was known (Fig. 2D), we re-fitted ELC models (and selected the best model via AIC) while setting *λ* to its true profile over time. To construct diversification scenarios congruent to the fitted models, but with the correct *μ* (e.g. Fig. 4C), we used the castor function `simulate_deterministic_hbd`.

Extant timetrees of 121 eukaryotic taxa were obtained from the literature; many of these timetrees had been previously collected from the literature by Henao Diaz *et al.* (32), and some were obtained by extracting sub-trees corresponding to recognized taxa (families or orders) from a large mammal tree (33) or avian tree (34). Taxon-specific sub-trees were only extracted if the tips associated with the taxon indeed formed a monophyletic clade in the original tree, based on the NCBI taxonomy (status December 3, 2020). The sampling fraction *ρ* of each tree was calculated based on published estimates of the taxon’s total number of extant species; tree sources and literature references are given in Supplemental Data File 1. To each tree, we fit all ELC models while fixing *ρ* and chose the best supported ELC model according to the AIC (35).

Statistical tests for model adequacy were performed using parametric bootstrapping (26, 27), as follows. For a given tree and a given fitted model to be evaluated, we simulated 1000 random trees from the model, fixing the number of tips and the root age to match those of the original tree. Simulations were performed using the castor function `generate_tree_hbd_reverse` (31). For every simulated tree, we calculated the Sackin statistic (21) (denoted *σ*), and then calculated the mean Sackin statistic across all simulated trees (denoted 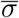). The statistical significance of the tree’s Sackin statistic (denoted *σ_o_*) was the fraction of simulated trees for which 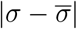 was greater than 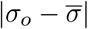. A similar approach was taken for the Colless statistic. To compare the distribution of node ages (times before present) of the fitted model to the tree, we used a Kolmogorov-Smirnov test. Specifically, for every simulated tree we calculated the empirical cumulative distribution function (CDF) of the node ages (denoted *F*), evaluated at the original tree’s node ages via linear interpolation, and then calculated the average of those CDFs, thus obtaining an estimate for the CDF of node ages expected under the model (denoted 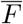). The Kolmogorov-Smirnov (KS) distance between a tree’s CDF *F* and 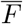, denoted 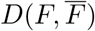, is the maximum distance between *F* and 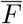 at any age. The statistical significance of the original tree’s KS distance 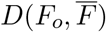 was calculated as the fraction of simulated trees for which 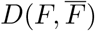 was larger than 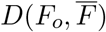. A similar approach was followed for comparing the model’s and tree’s distribution of edge lengths. The full array of statistical tests is implemented in the castor function model_adequacy_hbd (30). In all tests, the statistical significance threshold was set to 5%.

## Acknowledgements

We thank L. Harmon, J. Rolland, B. Neto-Bradley, K. Kaur, and L.F. Henao Diaz for comments on the manuscript. MWP was supported by a NSERC Discovery Grant. SL was supported by a startup grant by the University of Oregon.

## Code availability

R code for performing our analyses is available at: www.loucalab.com/archive/ZeroExtinction

## Author contributions

Both authors contributed equally to this work.

## Supplementary Materials

**Figure S1:**
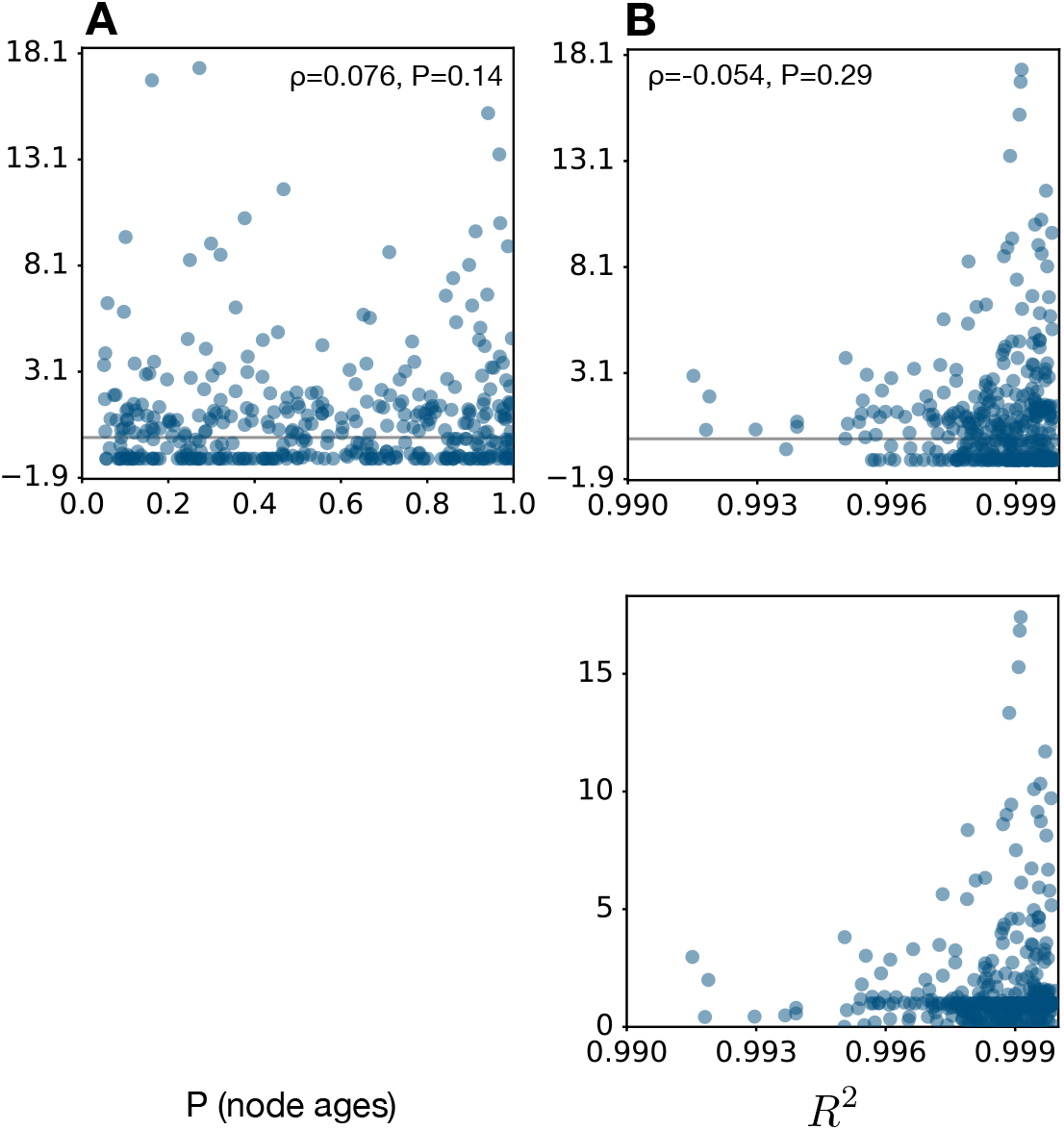
Relative estimation error vs. model adequacy (simulated trees). (A) Statistical significance of the Kolmogorov-Smirnov test for the distribution of node ages (horizontal axis) in 380 simulated trees under their best fitted constrained ELC models (one point per tree), compared to the relative error of the present-day extinction rate estimated under that model 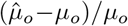, where *μ_o_* is the present-day extinction rate of the true diversification scenario used in the simulations). The Spearman’s rank correlation between the relative error and the Kolmogorov-Smirnov test significance, as well as the two-sided statistical significance of that correlation, are shown inside the figure. (B) Similar to A, but with the horizontal axis showing the fraction of variance (*R*^2^) in the true scenario’s dLTT captured by the fitted model’s dLTT. Observe that all fitted models nearly exactly reproduced the dLTT of the true scenario, which means that the fitted models indeed converged towards the true scenario’s congruence class (as expected).

**Figure S2:**
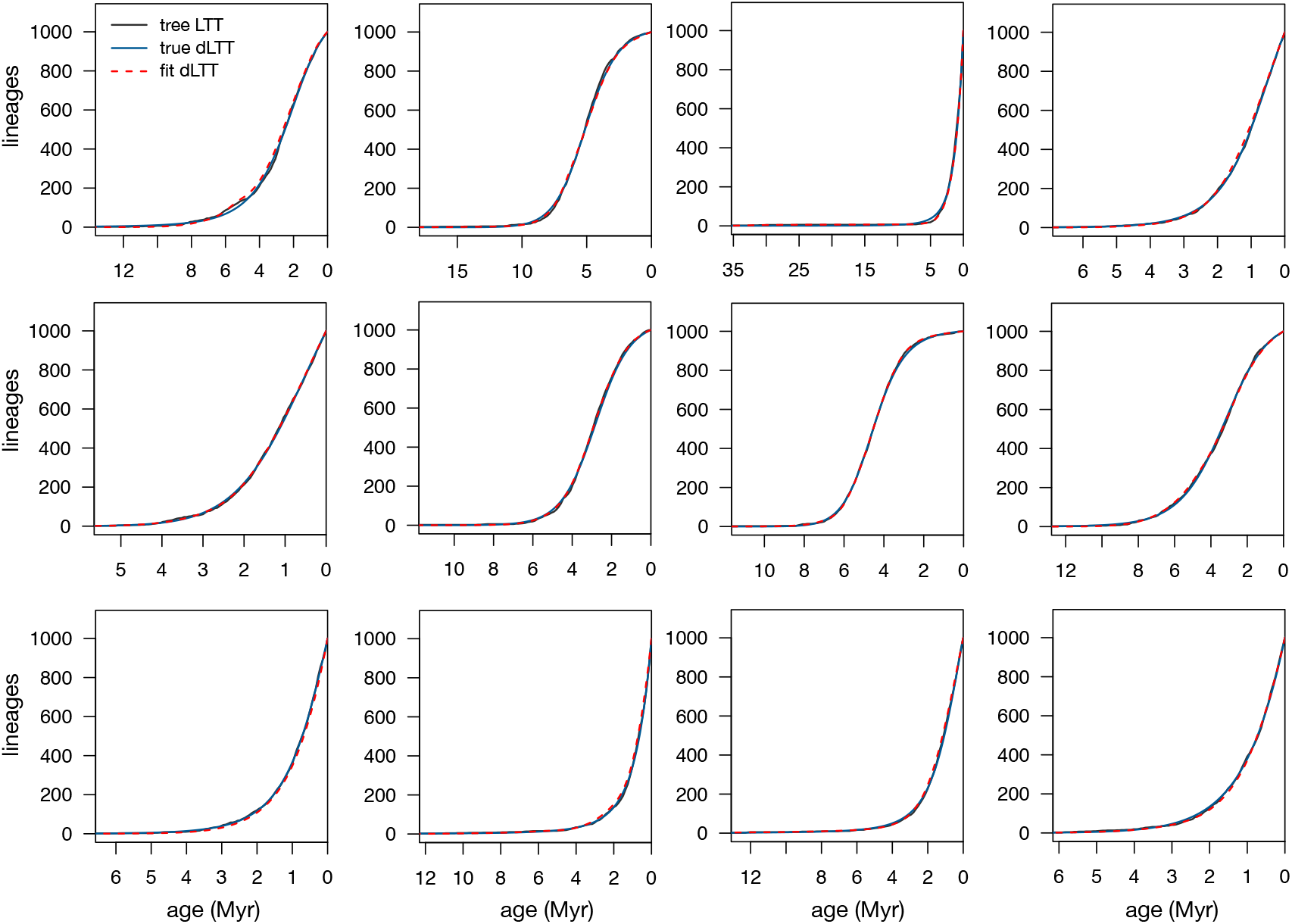
LTTs of trees vs. fitted models (simulated trees). Typical lineages-through-time (LTT) curves of timetrees simulated under various hypothetical diversification scenarios (continuous black curves), compared to the deterministic LTTs of the true generating diversification scenario (blue continuous curves) and the deterministic LTTs of the best fitted constrained ELC models (red dashed curves). In all examples shown the estimated present-day extinction rate 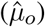 was zero.

**Figure S3:**
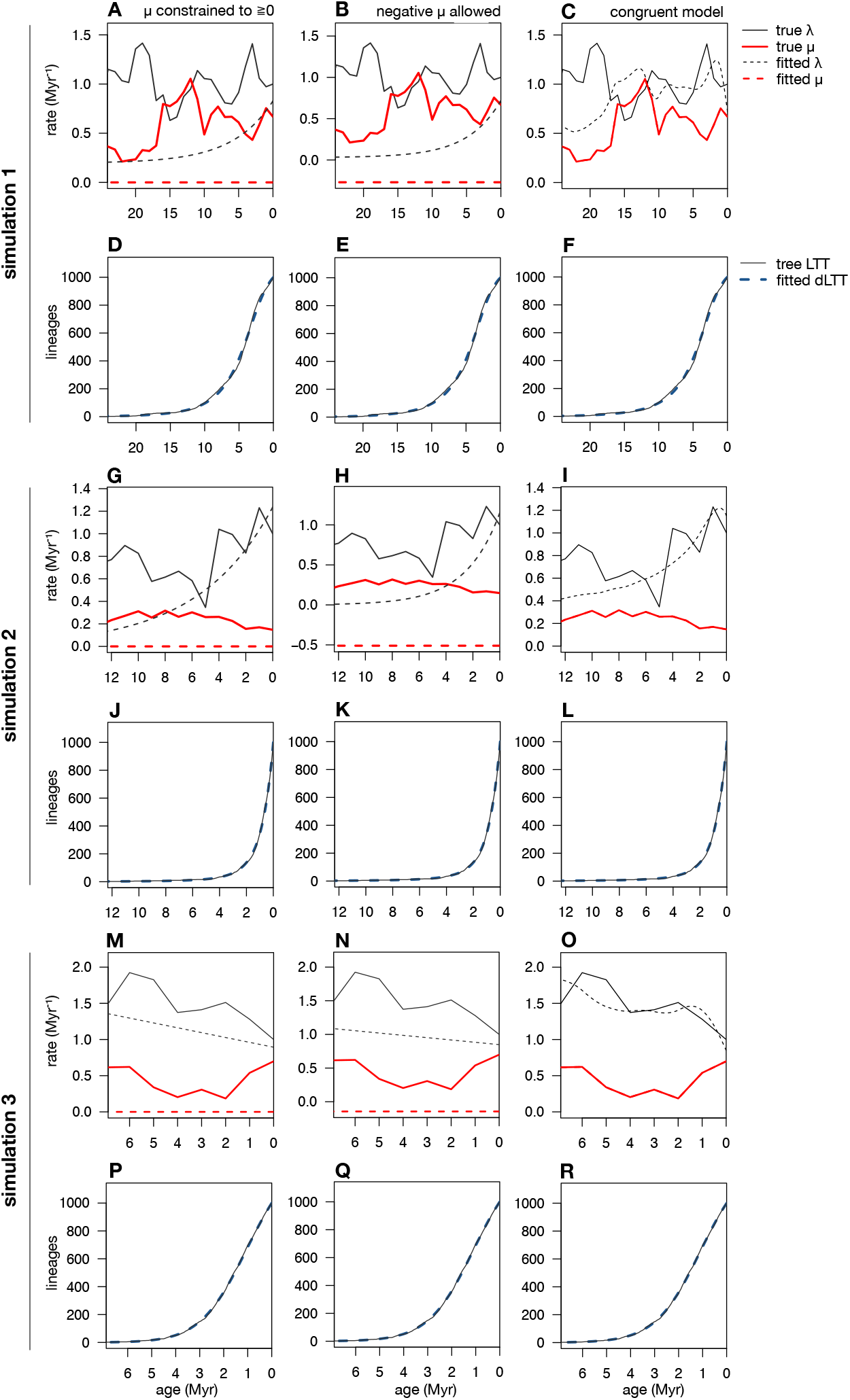
Fitted models are approximately congruent to the true diversification scenarios (simulated trees). Example ELC models fit (and selected according to AIC) to simulated timetrees with time-variable speciation rates (*λ*) and extinction rates (*μ*). (A) True *λ* and *μ* (continuous curves) compared to the *λ* and *μ* of the fit model (dashed curves), when *μ* is constrained to be non-negative. (B) True *λ* and *μ* (continuous curves) for the same tree as in A, compared to *λ* and *μ* (dashed curves) fitted when *μ* is allowed to be negative. (C) Model congruent to the model fitted in B, but exhibiting the correct extinction rate; observe that this model is close to the true diversification history, which means that the congruence class of the model fitted in B is close to the congruence class of the true diversification scenario. (D–F) Lineages-through-time (LTT) curve of the simulated tree (continuous curves) compared to the deterministic LTTs of the fitted models shown in A–C. The good match between the tree’s LTT and the fitted models’ deterministic LTTs is consistent with the conclusion that the fitted models are close to the congruence class of the true diversification scenario. (G–R) Similar to A–F, for 3 additional diversification scenarios.

**Figure S4:**
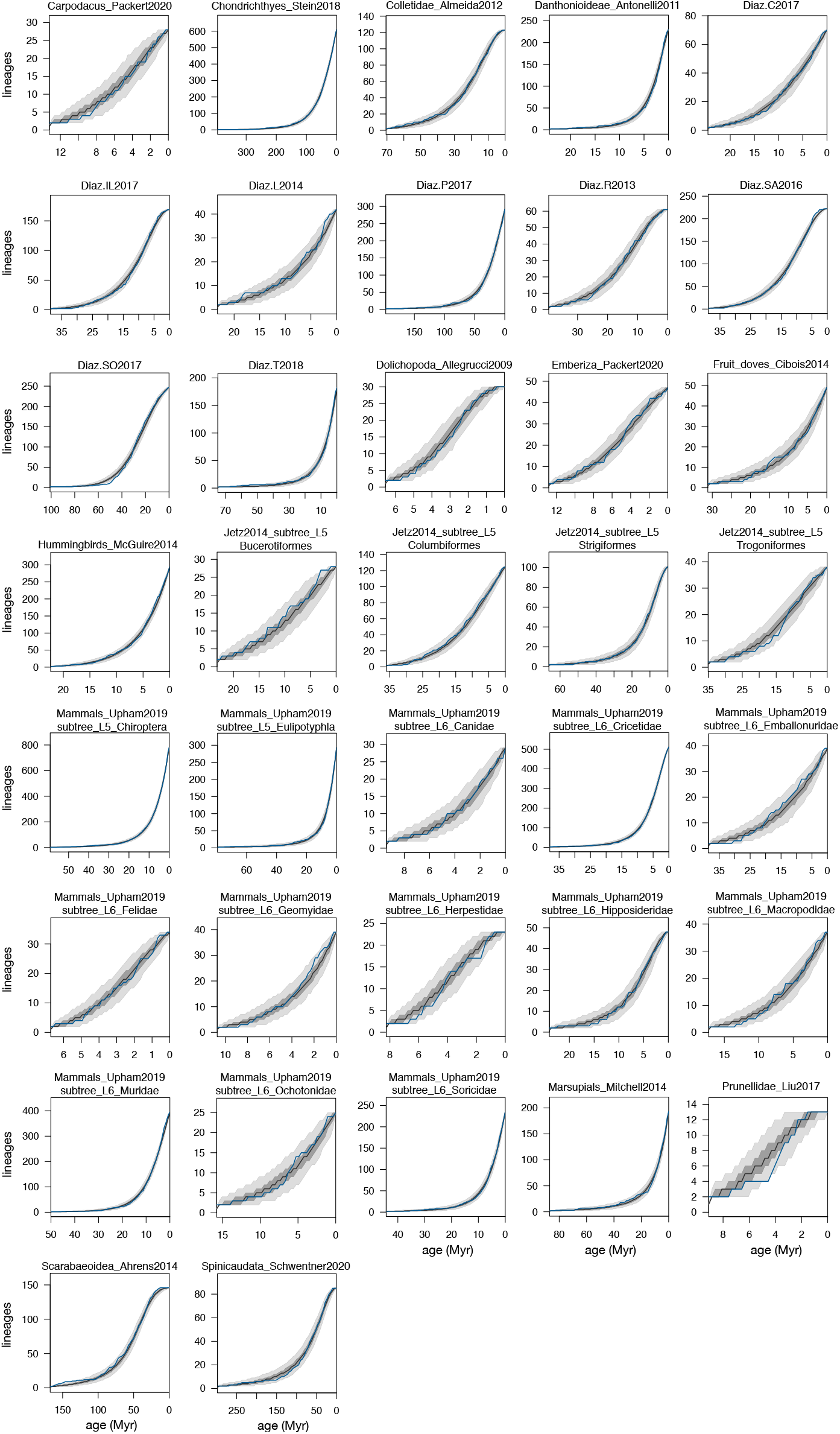
LTTs of trees vs. fitted models (empirical trees). Lineages-through-time (LTT) curves of the 37 empirical trees for which the best fitted ELC model adequately described the tree’s node distribution (based on a Kolmogorov-Smirnov test) and for which the predicted present-day extinction rate was zero (if constrained) or negative (if unconstrained). Blue curves show the LTTs of the trees, black curves show the median LTT generated by the fitted model, and dark and light shades show the 50% and 95% equal-tailed confidence intervals of LTTs generated by the fitted model. Note the large consistency between the tree LTTs and the LTTs generated by the fitted models.

**Figure S5:**
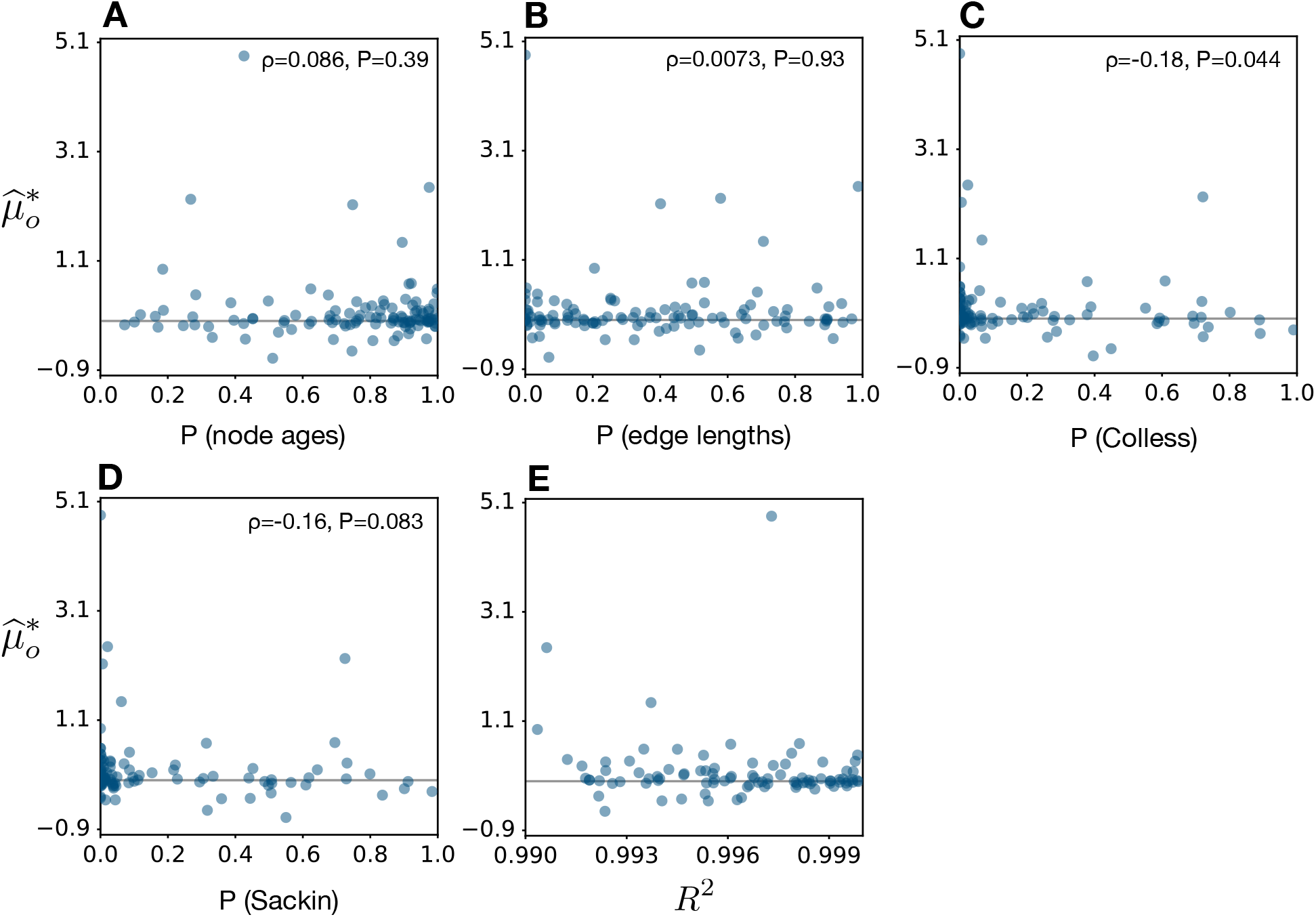
Extinction rate estimates vs. model adequacy (empirical trees). (A) Statistical significance of the Kolmogorov-Smirnov test for the distribution of node ages (horizontal axis) in empirical trees under their best fitted ELC models (one point per tree), compared to the present-day extinction rate estimated under that model (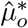, allowing for negative extinction rates). The Spearman’s rank correlation between 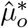 and the Kolmogorov-Smirnov test significance, as well as the two-sided statistical significance of that correlation, are shown inside the figure. (B) Similar to A, but for the distribution of edge lengths. (C) Statistical significance of each tree’s Colless statistic (horizontal axis) under the best fitted ELC model, compared to the corresponding 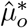 (one point per tree). (D) Similar to C, but for the Sackin statistic. (D) Fraction of variance in the tree’s LTT explained by the fitted model’s dLTT (*R*^2^), compared to the corresponding 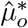 (one point per tree). None of the metrics shown in the horizontal axes (P-values and *R*^2^) were found to significantly positively correlate with 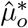, suggesting that model inadequacy unlikely drives the many negative (or zero, if constrained) extinction rate estimates.

## Notes

### Competing Interest Statement

The authors have declared no competing interest.

## References

[1] MacLeod N (2014) The geological extinction record: History, data, biases, and testing. Geological Society of America Special Papers 505:1–28.

[2] Marshall CR (2017) Five palaeobiological laws needed to understand the evolution of the living biota. Nature Ecology & Evolution 1(6):165.

[3] Pimm SL et al. (2014) The biodiversity of species and their rates of extinction, distribution, and protection. Science 344(6187):1246752.

[4] Nee S, Holmes EC, May RM, Harvey PH (1994) Extinction rates can be estimated from molecular phylogenies. Philosophical Transactions: Biological Sciences 344(1307):77–82.

[5] Losos JB, Schluter D (2000) Analysis of an evolutionary species–area relationship. Nature 408(6814):847–850.

[6] Turgeon J, Stoks R, Thum RA, Brown JM, McPeek MA (2005) Simultaneous quaternary radiations of three damselfly clades across the holarctic. The American Naturalist 165(4):E78–E107.

[7] Nee S (2006) Birth-death models in macroevolution. Annual Review of Ecology, Evolution, and Systematics 37(1):1–17.

[8] Purvis A (2008) Phylogenetic approaches to the study of extinction. Annual Review of Ecology, Evolution, and Systematics 39(1):301–319.

[9] Steeman ME et al. (2009) Radiation of extant cetaceans driven by restructuring of the oceans. Systematic Biology 58(6):573–585.

[10] Quental TB, Marshall CR (2010) Diversity dynamics: molecular phylogenies need the fossil record. Trends in Ecology & Evolution 25(8):434–441.

[11] Herrera JP (2017) Primate diversification inferred from phylogenies and fossils. Evolution 71(12):2845–2857.

[12] Rabosky DL (2010) Extinction rates should not be estimated from molecular phylogenies. Evolution 64(6):1816–1824.

[13] Morlon H, Parsons TL, Plotkin JB (2011) Reconciling molecular phylogenies with the fossil record. Proceedings of the National Academy of Sciences 108(39):16327–16332.

[14] Beaulieu JM, O’Meara BC (2015) Extinction can be estimated from moderately sized molecular phylogenies. Evolution 69(4):1036–1043.

[15] Rabosky DL (2016) Challenges in the estimation of extinction from molecular phylogenies: a response to beaulieu and o’meara. Evolution 70(1):218–228.

[16] Foote M (2003) Origination and extinction through the phanerozoic: a new approach. The Journal of Geology 111(2):125–148.

[17] Louca S, Pennell MW (2020) Extant timetrees are consistent with a myriad of diversification histories. Nature 580:502–505.

[18] Pagel M (2020) Evolutionary trees can’t reveal speciation and extinction rates. Nature 580:461–462.

[19] Stadler T, Steel M (2019) Swapping birth and death: Symmetries and transformations in phylodynamic models. Systematic Biology 68(5):852–858.

[20] McGuire JA et al. (2014) Molecular phylogenetics and the diversification of hummingbirds. Current Biology 24(8):910–916.

[21] Sackin MJ (1972) ‘good’ and ‘bad’ phenograms. Systematic Biology 21(2):225–226.

[22] Shao KT (1990) Tree balance. Systematic Biology 39(3):266–276.

[23] Uhlenbeck GE, Ornstein LS (1930) On the theory of the Brownian motion. Physical Review 36:823–841.

[24] Alroy J (2008) Dynamics of origination and extinction in the marine fossil record. Proceedings of the National Academy of Sciences 105(Supplement 1):11536–11542.

[25] Rabosky DL, Lovette IJ (2008) Explosive evolutionary radiations: decreasing speciation or increasing extinction through time? Evolution: International Journal of Organic Evolution 62(8):1866–1875.

[26] Brown JM, Thomson RC (2018) Evaluating model performance in evolutionary biology. Annual Review of Ecology, Evolution, and Systematics 49(1):95–114.

[27] Schwery O, O’Meara BC (2020) Boskr–testing adequacy of diversification models using tree shape. bioRxiv.

[28] Nee S, May RM, Harvey PH (1994) The reconstructed evolutionary process. Philosophical Transactions of the Royal Society of London B: Biological Sciences 344(1309):305–311.

[29] Lambert A, Stadler T (2013) Birth–death models and coalescent point processes: The shape and probability of reconstructed phylogenies. Theoretical Population Biology 90:113–128.

[30] Louca S, Doebeli M (2018) Efficient comparative phylogenetics on large trees. Bioinformatics 34(6):1053–1055.

[31] Louca S (2020) Simulating trees with millions of species. Bioinformatics 36(9):2907–2908.

[32] Henao Diaz LF, Harmon LJ, Sugawara MTC, Miller ET, Pennell MW (2019) Macroevolutionary diversification rates show time dependency. Proceedings of the National Academy of Sciences 116(15):7403.

[33] Upham NS, Esselstyn JA, Jetz W (2019) Inferring the mammal tree: Species-level sets of phylogenies for questions in ecology, evolution, and conservation. PLoS biology 17(12):e3000494.

[34] Jetz W et al. (2014) Global distribution and conservation of evolutionary distinctness in birds. Current Biology 24(9):919–930.

[35] Akaike H (1981) Likelihood of a model and information criteria. Journal of Econometrics 16(1):3–14.

